# Mapping Structural and Dynamic Divergence Across the MBOAT family

**DOI:** 10.1101/2023.12.18.572254

**Authors:** T. Bertie Ansell, Megan Healy, Claire E. Coupland, Mark S. P. Sansom, Christian Siebold

**Affiliations:** Department of Biochemistry, South Parks Road, Oxford, OX1 3QU, UK; Division of Structural Biology, Wellcome Centre for Human Genetics, Roosevelt Drive, Oxford, OX3 7BN, UK; Molecular Medicine Program, The Hospital for Sick Children, 686 Bay Street, Toronto, M5G 0A4, Canada; Division of CryoEM and Bioimaging, SSRL, SLAC National Accelerator Laboratory, Menlo Park, CA 94025, USA; Department of Biology, Stanford University, Stanford, CA 94305, USA

**Keywords:** Simulation, molecular dynamics, MBOAT, enzyme, catalysis, membrane protein, lipids, bilayer

## Abstract

Membrane Bound *O*-acyltransferases (MBOATs) are membrane embedded enzymes which catalyse acyl chain transfer to a diverse group of substrates including lipids, small-molecules and proteins. Recent MBOAT structures reveal a conserved structural core, despite wide-ranging functional specificity across both prokaryotes and eukaryotes. The structural basis of catalytic specificity, regulation and interactions with the surrounding environment remain uncertain, hindering effective therapeutic targeting. Here, we combine comparative molecular dynamics (MD) simulations with bioinformatics to assess molecular and interactional divergence across the family. In simulations, MBOATs differentially distort the bilayer depending on their substrate type. Additionally, we identify specific lipid binding sites surrounding MBOAT reactant gates into the surrounding membrane. We use bioinformatics to reveal a conserved role for re-entrant loop-2 in stabilisation of the MBOAT fold and identify a key hydrogen bond involved in DGAT1 dimerisation. Finally, we predict differences in MBOAT core solvation and water gating properties across the family. These data are pertinent to the design of MBOAT specific inhibitors that encompass dynamic information within cellular mimetic environments.

## Introduction

Membrane bound *O*-acyltransferases (MBOATs) are a family of membrane embedded enzymes found across prokaryotes and eukaryotes. The MBOAT family can be subdivided into two broad subfamilies dependent on whether an acyl chain is transferred from acyl-coenzyme A (acyl-CoA) onto either a protein or small-molecule acceptor^1^. For example, small-molecule acylating MBOATs include acyl-CoA:cholesterol acyltransferase (ACAT1), diacylglycerol acyltransferase (DGAT1) and lysophospholipid acyltransferases (LPCAT) which catalyse the acylation of cholesterol, diacylglycerol (DAG) and lysophospholipids respectively^2–8^. By contrast, protein acylating MBOATs include the morphogen acylating enzymes Hedgehog acyltransferase (HHAT) and Porcupine (PORCN) and the ghrelin *O-*acyltransferase (GOAT)^9–12^. Also included within this subfamily is the prokaryotic teichoic acid D-alanyltransferase, DltB, which is functionally distinct and catalyses D-alanylation of acids within the peptidoglycan layer of the cell-wall^13^.

The first MBOAT family structure, of DltB, was determined by X-ray crystallography^13^. The DltB structure revealed an archetypal MBOAT fold composed of a funnel of tilted transmembrane helices surrounding the catalytic reaction centre. More recently, there has been an explosion in MBOAT structures, facilitated by widespread application of cryogenic electron microscopy (cryo-EM)^14^. These structures unveil a conserved MBOAT fold comprised of eight transmembrane helices (hereafter TM1’-TM8’ where superscript’ denotes core numbering) positioned between structurally divergent N- and C-terminal helices^1^ (Fig. 1A). Additionally, two re-entrant loops between TM3’-TM4’ (re-entrant loop-1) and TM5’-TM6’ (re-entrant loop-2) contribute to the MBOAT core. Most MBOATs were structurally characterised as monomers with the exception of ACAT1, DGAT1 and LPCAT3 which were characterised as dimers (or dimers of dimers)^2–7^. For DGAT1 and ACAT1 the dimeric interface is positioned surrounding re-entrant loop-2 (Fig. 1A).

**Figure 1:**
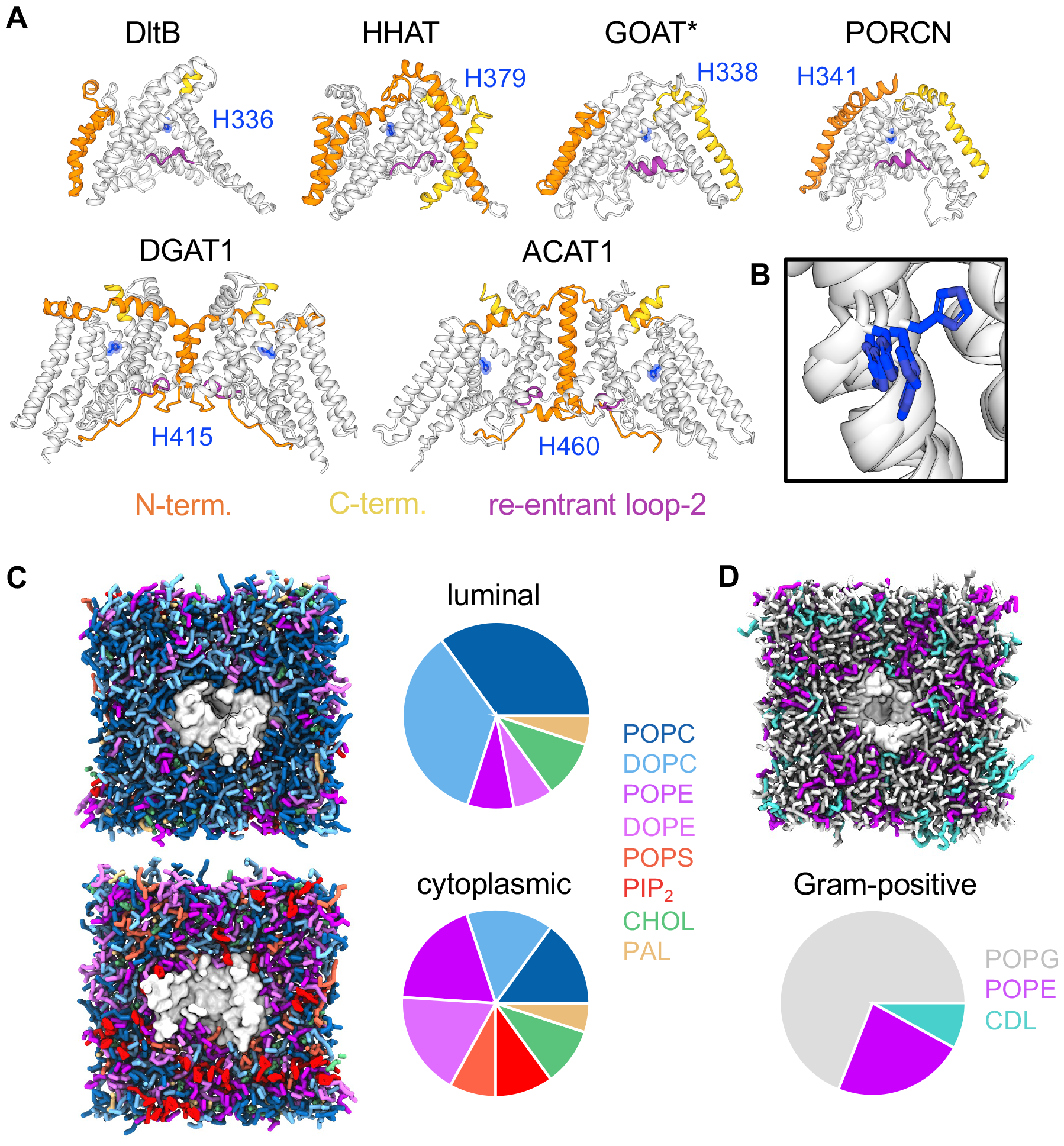
MBOAT family structures and membranes. **A)** Structures and models used in simulations of membrane bound *O-*acyltransferase (MBOAT) family members DltB (PDB: 6BUG^13^), Hedgehog acyltransferase (HHAT, PDB: 7Q1U^9^), ghrelin *O-*acyltransferase (GOAT, Uniprot: Q96T53), Porcupine (PORCN, PDB: 7URA^11^), diacylglycerol acyltransferase (DGAT1, PDB: 6VP0^5^) and acyl-CoA:cholesterol acyltransferase (ACAT1, PDB: 6P2P^2^) coloured as follows: the MBOAT core (white), N-terminus (orange), C-terminus (yellow), re-entrant loop-2 (purple) and the conserved catalytic histidine (blue). The GOAT model (marked *) was derived from the AlphaFold Protein Structure Database^36^. All other MBOATs were determined experimentally. **B)** Overlay of the position of the conserved catalytic histidine on TM6’ across MBOATs shown in **A**. Coarse-grained (CG) representations of **C)** HHAT embedded in an asymmetric ER mimetic bilayer composed of POPC (blue), DOPC (light blue), POPE (purple), DOPE (pink), POPS (orange), PIP_2_ (red), cholesterol (green) and palmitate (ochre) and **D)** DltB embedded in a Gram-positive like membrane composed of POPG (grey), POPE purple) and cardiolipin (CDL, teal). Pie charts indicate the relative lipid % compositions.

MBOAT catalysis is thought to be mechanised by two key residues; an invariant His residue and either Asn, Asp or His. Despite low sequence identity across the family, the position of these residues is structurally conserved on TM6’ (His) (Fig. 1B) and re-entrant loop-2 (Asn/Asp/His)^15^. The MBOAT fold positions these proposed catalytic residues approximately at the bilayer midplane like pincers on either side of the acyl-CoA thioester bond^2–11,13^. Hence, MBOATs contain a central enzymatic cavity for acyl-CoA and substrate engagement however little is known about how properties of this cavity differ to promote reaction specificity across the MBOAT family.

Several studies opened structural windows into the role of lipids in MBOAT function. In cryo-EM structures of ACAT1, ACAT2 and DGAT1 a membrane exposed cavity (termed the lateral gate) connects the reaction centre to the surrounding membrane^2–6,16^. Furthermore, cholesterol was successfully docked into this cavity of ACAT1^3^ and modelled at a similar position within a subsequent structure of ACAT2^16^. Membrane bending/deformations are observed within the bilayer spanning regions of ACAT1 and HHAT cryo-EM structures^4,9^. Complementary molecular dynamics (MD) simulations of HHAT further stipulate formation of membrane deformations, proposed to reduce the energetic cost of cross membrane catalytic transfer^9^. The importance of these observations is not to be understated given the implicit role of lipids and lipid-like substrate in MBOAT catalysis but remains unexplored across other MBOATs.

Detailed structural interpretation (e.g. side-chain rearrangements or resolved water molecules) may be assisted by computational analysis such as MD simulations or bioinformatics, particularly at resolutions which are not yet routinely reached for membrane protein cryo-EM structures^17^. MD simulations can help to shed lights on dynamic aspects which may be mechanistically important for protein function. These include protein gating, solvent/membrane accessibility of potentially druggable pockets or protein-lipids interactions^18,19^.

Here, we performed comparative MD simulations and bioinformatic analysis of six MBOATs across both subfamilies. We identify multiple distinct hallmarks of subfamily specialisation including differences in the extent and position of membrane deformations, reaction centre solvation and protein gating. We combine simulation and bioinformatic analysis to elucidate conserved roles for fold stabilisation at re-entrant loop-2 and dimeric tail-swap in DGAT1. These data provide a comprehensive computational platform for protein specific divergence across the MBOAT family which may be exploited in future experimental studies and/or for tailored pharmacological targeting.

## Results

Given the role of lipids and lipid-derivatives in MBOAT catalysis, we sought to assess protein accommodation within bilayers representative of their native environment. We performed CG simulations of six MBOATs (DltB, HHAT, PORCN, GOAT, DGAT1 and ACAT1) (Fig. 1), within membranes designed to mimic the endoplasmic reticulum (Fig. 1C) or Gram-positive cellular membrane (for DltB) (Fig. 1D).

### MBOAT subfamilies differentially alter membrane thickness

Visualisation of CG trajectories revealed marked regions of membrane deformation around a subset of MBOATs. Given the unusual trapezoid tertiary structure of MBOATs^1^, we sought to assess whether the extent and location of deformations was conserved across the family. For protein acylating MBOATs (HHAT, GOAT, PORCN and DltB) the membrane width was decreased by 1.5-1.8 nm between the most extreme regions of deformation along the z axis compared to the extended distances from the protein (global deformation) (Fig. 2). On the luminal/extracellular (EC) face the most pronounced region of deformation was conserved between HHAT, PORCN and DltB and localised to the proposed position of a luminal product exit gate between TM2’ and TM6’ of the MBOAT core (where superscript’ is used in accordance with standardised numbering of core helices across the family)^1^ (Fig. 2A, Supplementary Fig. 1, black arrow). This deformation funnels inwards towards the conserved catalytic histidine on TM6’. For GOAT, the maximal luminal deformation occurred in proximity to the N-terminus. On the cytoplasmic/intracellular (IC) face, the region of reduced membrane width coincided with re-entrant loop-2 for all protein MBOATs (Fig. 2A, Supplementary Fig. 1, black asterisks), as previously noted in proximity to the heme binding site on HHAT^9^.

**Figure 2:**
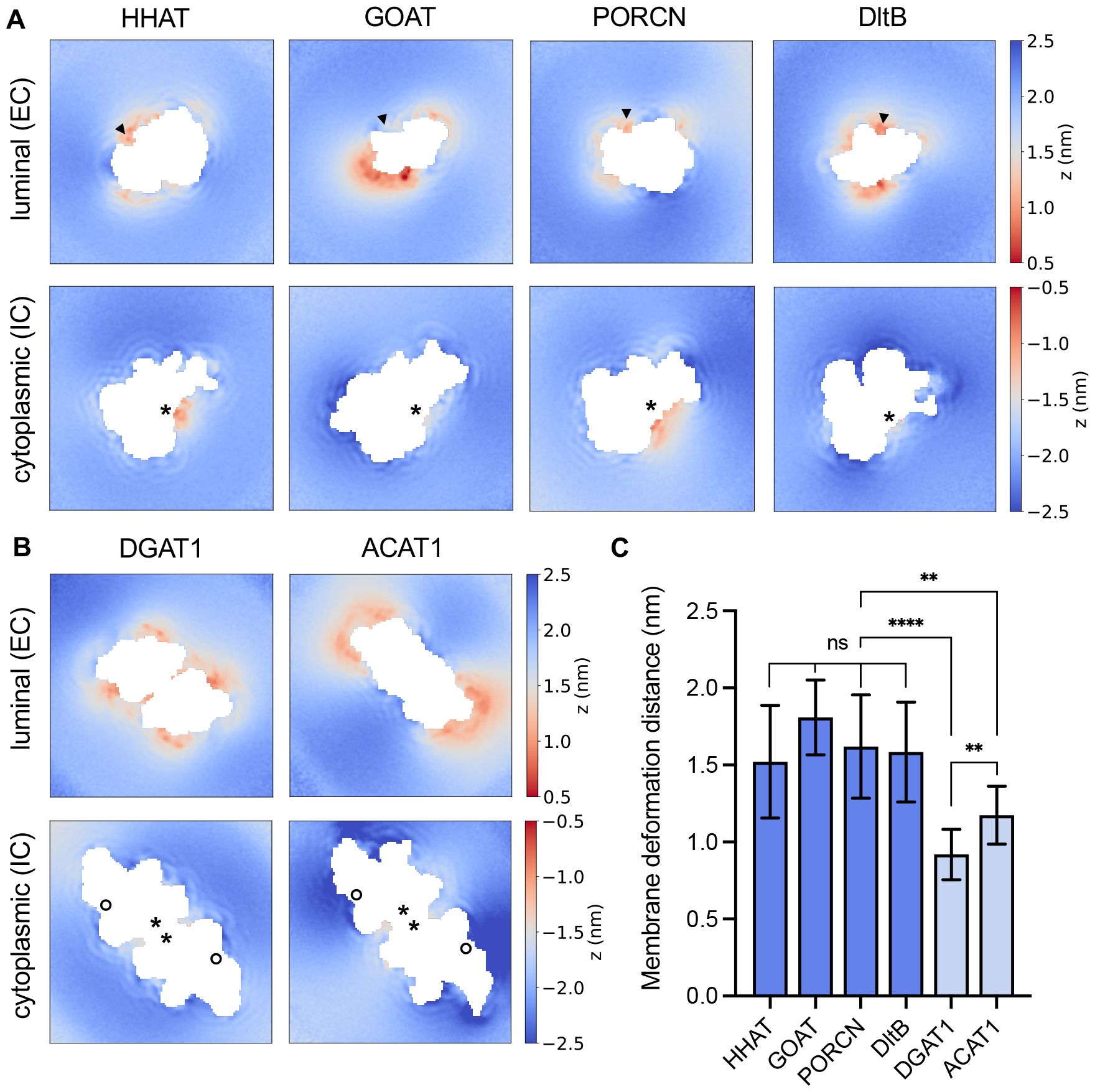
MBOATs induce membrane deformation. Time averaged z axial position of membrane phosphate beads across 10 x 15 μs CG simulations of **A)** protein acylating MBOATs (HHAT, GOAT, PORCN and DltB) and **B)** small-molecule MBOATs (DGAT1 and ACAT1 dimers). The bilayer midplane (z=0 nm) was defined as the mean phosphate z position. The position of phosphates within the luminal/extracellular (EC) or cytoplasmic/intracellular (IC) leaflets were normalised to the bilayer midplane and plotted as a binned 2D array surrounding proteins. Black arrows indicate the position of the luminal gate between TM2’ and TM6’. Asterisks mark the position of re-entrant loop-2 and the circle shows the position of the lateral gate. **C)** Bar plot of global membrane deformation, defined as the reduction in bilayer width between the most extreme regions of deformation surrounding each MBOAT compared to phosphate positions at extended distances from the protein (plotted as mean ± s.d. of phosphate bead positions). Statistical significance was determined by a Students unpaired t-test: not-significant (ns): *P* > 0.05, ^*^: *P* ≤ 0.05, ^**^: *P* ≤ 0.01, ^***^: *P* ≤ 0.001, ^****^: *P* ≤ 0.0001.

By contrast, small-molecule MBOATs ACAT1 and DGAT1 induced significantly less global membrane deformation (0.9-1.2 nm) than protein MBOATs, which mostly localised to the luminal face (Fig. 2B-C). Of note, we did not observe any deformation around the lateral gate (Fig. 2B, Supplementary Fig. 1-2A, black circle) and the ACAT1/DGAT1 dimer interface occludes the region of cytoplasmic deformation observed in protein MBOATs (black asterisks). Differential degrees of membrane deformation were also observed for lipid phosphate beads proximal to the protein compared to the most extreme regions of deformation (local deformation) between protein and small-molecule MBOATs (Supplementary Fig. 2B). Hence protein and small-molecule MBOAT subfamilies induce membrane thinning by different degrees, with protein acylating MBOATs substantially reducing (∼40-60%) the membrane width compared to at extended distances from the protein. Mechanistically, membrane thinning may reduce the energetic cost of substrate transfer across the bilayer and/or toward the catalytic core. Given protein MBOAT catalysis involves acyl/D-alanyl transfer across the membrane (rather than into the membrane as occurs for small-molecule MBOATs), conserved regions of localised membrane distortion may represent one hallmark of subfamily specialisation to facilitate reactant entry and release.

### Kinetics of specific lipid interactions surrounding key gating sites

Encouraged by our assessment of global changes in membrane thickness around MBOAT subfamilies, we assessed whether there were differences in specific protein-lipid interactions between MBOATs. We used PyLipID^20^ to calculate protein-lipid binding sites and their associated kinetics across the MBOAT family (Fig. 3). A number of lipid binding sites were observed for each MBOAT; therefore, we chose to focus on comparing sites surrounding regions pertinent to proposed catalytic mechanisms. For protein MBOATs we observe a phospholipid binding site between the luminal gate helices on TM2’ and TM6’ (Fig. 3A, Binding Site-1). For PORCN and GOAT, the phospholipid headgroups of POPC, DOPC, POPE and DOPE fold over the luminal surface of TM6’ towards the catalytic headgroup with relatively little kinetic specificity between PC and PE lipids (Fig. 3A, residence time plot). Lipids also bind to Binding Site-1 on DltB, with a marked increase in residence time for cardiolipin compared to POPG and POPE. This site has been previously investigated on HHAT whereby the DOPC headgroup also arrowed towards H379 in CG simulations. During extended atomistic simulations one tail of DOPC occupied the reaction centre, acting as a product mimetic to open the luminal gate^9^. Hence, for protein-MBOATs Binding Site-1 may represent a conserved phospholipid binding site whereby membrane lipids occupy the luminal gate periphery until they are displaced by the exiting acylated product. Notably, for DltB we observed a second prominent lipid binding site directly below the luminal gate (Fig. 3B, Binding Site-2). This site has a residence time of 15 μs for POPG, POPE and cardiolipin which appeared to be driven by shape complementarity within a cavity between TM2’/TM6’ helices rather than headgroup specificity. The role of this prominent binding site remains to be investigated but was also seen for PIP_2_ in CG simulations of HHAT^21^.

**Figure 3:**
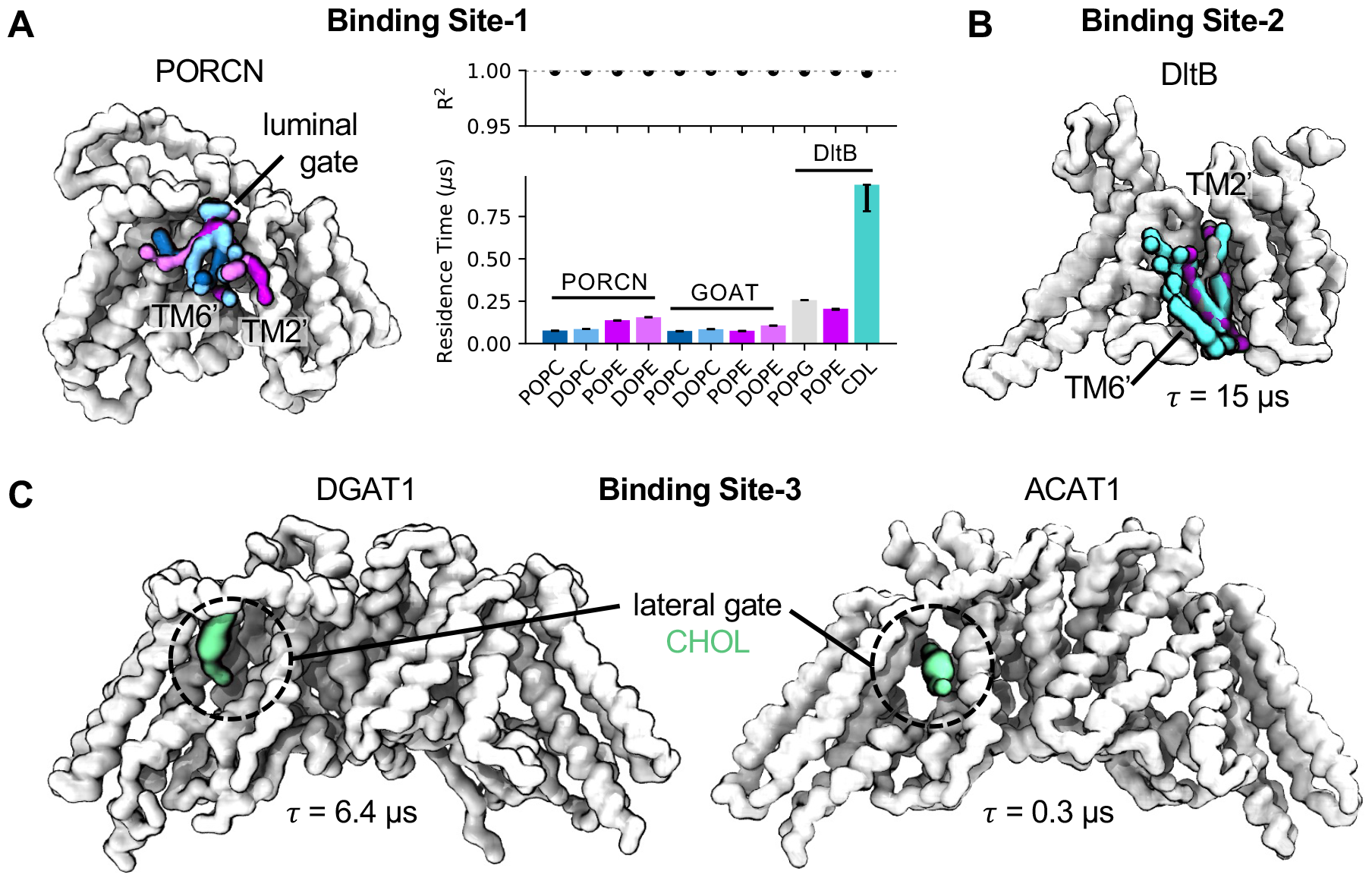
Lipid interactions at the luminal and lateral gates. Protein-lipid interactions surrounding the luminal gate of protein MBOATs (**A-B**) and lateral gate of small-molecule MBOATs (**C**). Top ranked lipid binding poses and lipid residence times were identified using PyLipID^20^ from 10 x 15 μs CG simulations of each MBOAT. **A)** Binding Site-1: POPC (blue), DOPC (light blue), POPE (purple) and DOPE (pink) bound to the luminal gate of PORCN (white). MBOAT core helices are numbered with superscript’ as defined in ^1^. A residence time (*τ*) comparison plot for lipids bound to PORCN, GOAT and DltB at Binding Site-1 is shown, adapted from LipIDens^21^ outputs. R^2^ values for the biexponential fit of *k*_*off*_ values (where *τ* = 1/*k*_*off*_) are indicated and asymmetric error bars correspond to residence times for *k*_*off*_ values obtained via bootstrapping to the same data. **B)** Binding Site-2: Cardiolipin (CDL, teal), POPG (grey) and POPE (purple) bound to a site on DltB situated directly below the luminal gate with a prolonged residence time (*τ* = 15 μs). **C)** Binding Site-3: Cholesterol (CHOL, green) within the lateral gate cavity of DGAT1 and ACAT1.

Next, we investigated lipid binding to the lateral gate of small-molecule MBOATs. Specifically, we sought to assess whether cholesterol (present in the ER mimetic membrane) bound to this site, given its role in ACAT1 catalysis and the similarity in size to DAG which functions in DGAT1 catalysis. We observe a cholesterol binding site within the lateral gate tunnel of both DGAT1 and ACAT1 in CG simulations (Fig. 3C, Binding Site-3). For DGAT1 the top ranked cholesterol pose was orientated with the ROH bead (equivalent to the 3β-hydroxy group) facing the gate periphery. Reassuringly, the cholesterol pose orientation in ACAT1 was aligned with the tunnel (i.e. parallel to the bilayer midplane) and the ROH bead faced inwards towards the catalytic cavity. This is in line with previous docking predictions for cholesterol within the ACAT1 lateral gate and the proposed catalytic mechanism^2^, albeit with a reduced residence time compared to Binding Site-3 on DGAT1.

### Conserved role for re-entrant loop-2 in stabilisation of the MBOAT fold

We performed bioinformatic analysis of MBOATs by mapping multiple-sequence alignments onto the protein structures (Fig. 4). We note a region of residue conservation surrounding re-entrant loop-2 across all MBOATs. This is intriguing since, re-entrant loop-2 forms the heme-b binding site for HHAT but the cysteine coordinating residue (C324) is not conserved in other MBOATs^9,10^ (Fig. 4A). Instead, MBOATs appear to have evolved distinct mechanisms of stabilising the MBOAT fold at this site. For GOAT and PORCN a conserved salt-bridge connects re-entrant loop-2 to the tilted TM5’ helix (Fig. 4B-C). In atomistic MD simulations of GOAT, K252 and E294 form a salt-bridge for 99.5 ± 0.4% of the total simulation time. In PORCN simulations, we observe rearrangement of H252, compared to the structural position, to form a salt-bridge with E293 for 52.3 ± 0.4% of one trajectory (Fig. 4B-C). For DltB, *π*-*π* stacking interactions are observed between conserved phenylalanine residues (F247-F276) at equivalent positions to charged residues in GOAT/PORCN (Fig. 4D). For DGAT1 and ACAT1 this site forms the dimeric interface (Supplementary Fig. 3). Hence, MBOATs have evolved distinct mechanisms of stabilising the MBOAT fold, including between subfamily members. This further supports a role for the HHAT heme-b in protein stabilisation rather that catalysis, in line with C324 mutational studies which disrupt HHAT folding^9^. It is intriguing that all these interactions (heme-b binding, salt-bridges, *π* - *π* interactions and dimer formation) are theoretically reversible and could represent different mechanisms of regulating MBOAT enzymatic activity under distinct cellular contexts.

**Figure 4:**
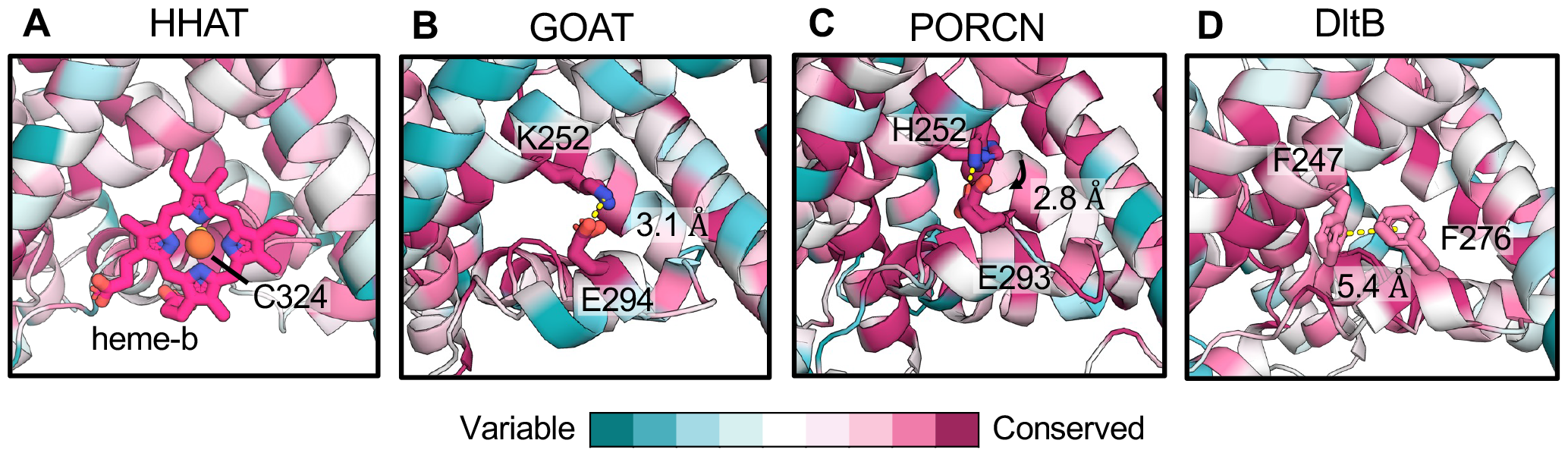
Divergent mechanisms of MBOAT fold stabilisation. Per residue sequence conservation mapped onto the structures/models of **A)** HHAT^9^, **B)** GOAT^36^, **C)** PORCN^11^ and **D)** DltB^13^. Bioinformatic analysis was derived from multiple-sequence alignments (MSAs) of individual proteins and mapped onto structures using ConSurf^37^. Conserved residues which coordinate the heme-b (HHAT), form salt-bridges (GOAT, PORCN) or *π*-*π* stacking interactions (DltB) are shown as sticks at the start and end of 200 ns atomistic simulations, with interaction distances labelled (**B-D**).

### A single hydrogen bond stabilises DGAT1 tail swap

In addition to the involvement of re-entrant loop-2 in DGAT1 dimer formation, we noticed a highly conserved hydrogen bond (H69/T260) between the N-terminal tail of one DGAT1 monomer and the cytosolic face of the neighbouring subunit (Fig. 5A-B). Unlike ACAT1, DGAT1 engages in N-terminal tail swap, suggested to be involved in regulation of the disordered N-terminal inhibitory domain^22^. Intrigued by the high conservation of H69 and T260 within an area of overall low sequence conservation, we performed atomistic simulations of DGAT1 to better assess protein dynamics. We calculated the root mean square deviation (RMSD) of the N-terminal tail Cα atoms and plotted against the prevalence of hydrogen bond formation between H69/T260. In two simulations (replicates 2 and 4) the H69-T260 hydrogen bond is broken, which correlated with dissociation of the N-terminal tail of DGAT1 from the adjacent subunit, indicated by a rapid increase in tail RMSD (Fig. 5C). When the H69/T260 interaction was stable (replicates 1, 3 and 5) the N-terminal tail remained bound. Hence both bioinformatic and simulation analyses suggest hydrogen bond formation between H69 and T260 is the dominant stabilising interaction within the DGAT1 N-termini. These data spotlight a specific molecular interaction that could be pharmacologically targeted for regulation of DGAT dimers compared to other MBOATs.

**Figure 5:**
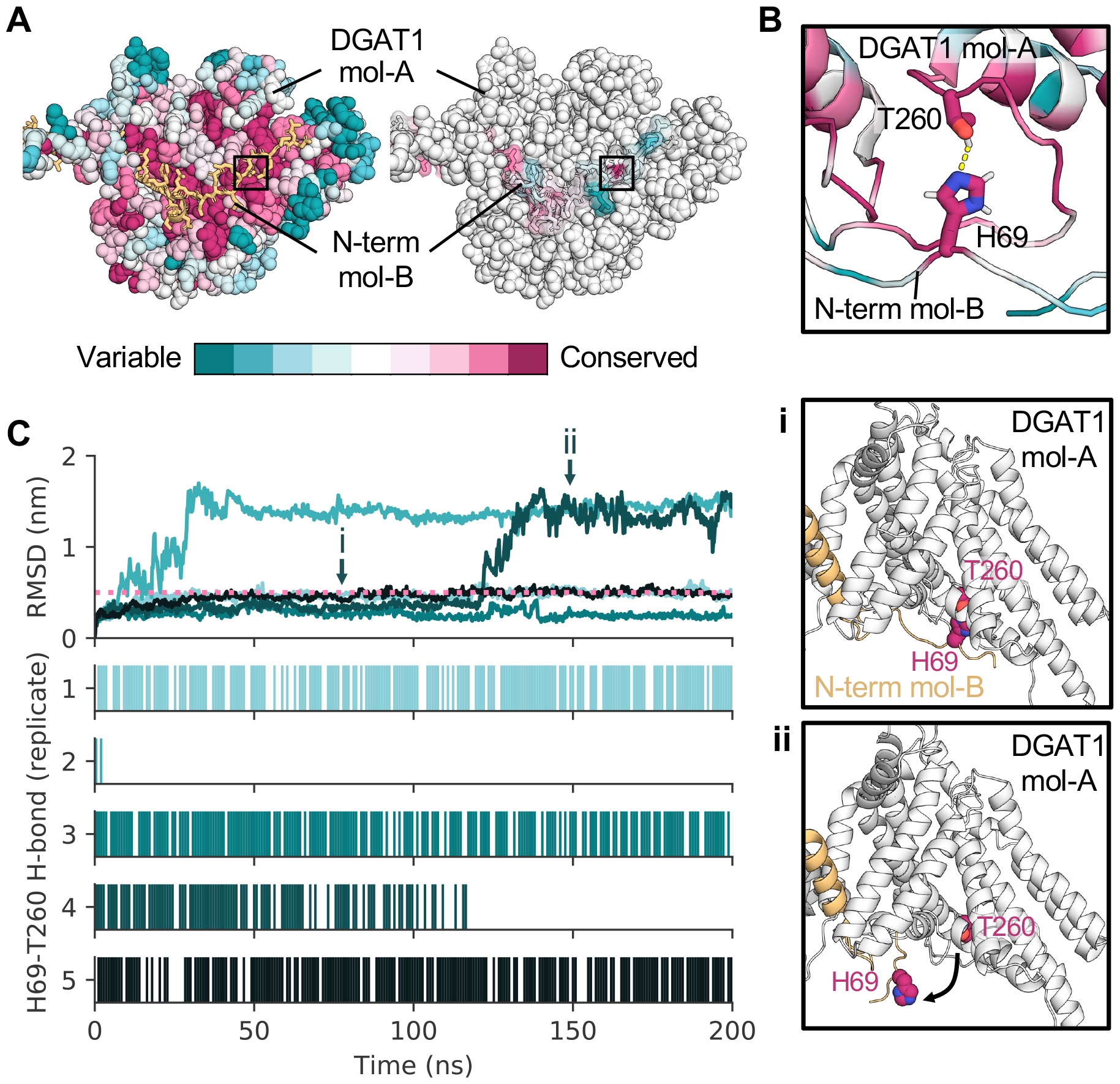
DGAT1 tail exchange is stabilised by a conserved hydrogen bond. A) Residue conservation mapped onto DGAT1 viewed from the cytosolic surface. For clarity one DGAT1 monomer is shown as spheres (mol-A) and the tail swapped N-terminus of the neighbouring DGAT1 monomer is shown as sticks (mol-B). A conserved hydrogen bond between H69 and T260 is boxed. **B)** Close up of the interaction between H69 and T260 coloured by sequence conservation. **C)** Root mean square deviation (RMSD) of DGAT1 N-term. Cα atoms (residues E65-R86) across 5 x 200 ns atomistic simulations. Replicates are coloured individually. The pink dashed line indicates the RMSD threshold below which the N-term. remains stabilised. The hydrogen bond prevalence (calculated using MDAnalysis^38^) is plotted for each replicate. Snapshots of tail stabilisation by the hydrogen bond (i) or after the interaction is broken (ii) are boxed.

### Water occupies the MBOAT reaction centre

We calculated the time averaged water number density across atomistic simulations of MBOATs to assess how solvent may affect more mechanistic aspects of enzyme catalysis. All MBOATs show a similar pattern of cavity solvation funnelling from the luminal/extracellular face towards the reaction centre (Fig. 6A, dashed circle). Hence, both conserved catalytic residues are likely hydrated in the absence of bound substrates. On the cytosolic/intracellular surface, re-entrant loop-2 is hydrated and situated at the membrane-solvent interface, consistent with our assessment of re-entrant loop-2 as a conserved region of membrane deformation (Fig. 2). Solvation of the MBOAT core may be a second mechanism (besides membrane deformation) of reducing the hydrophobic barrier for substrate transport into the centre of the membrane, and/or optimising conditions for efficient reaction centre catalysis.

**Figure 6:**
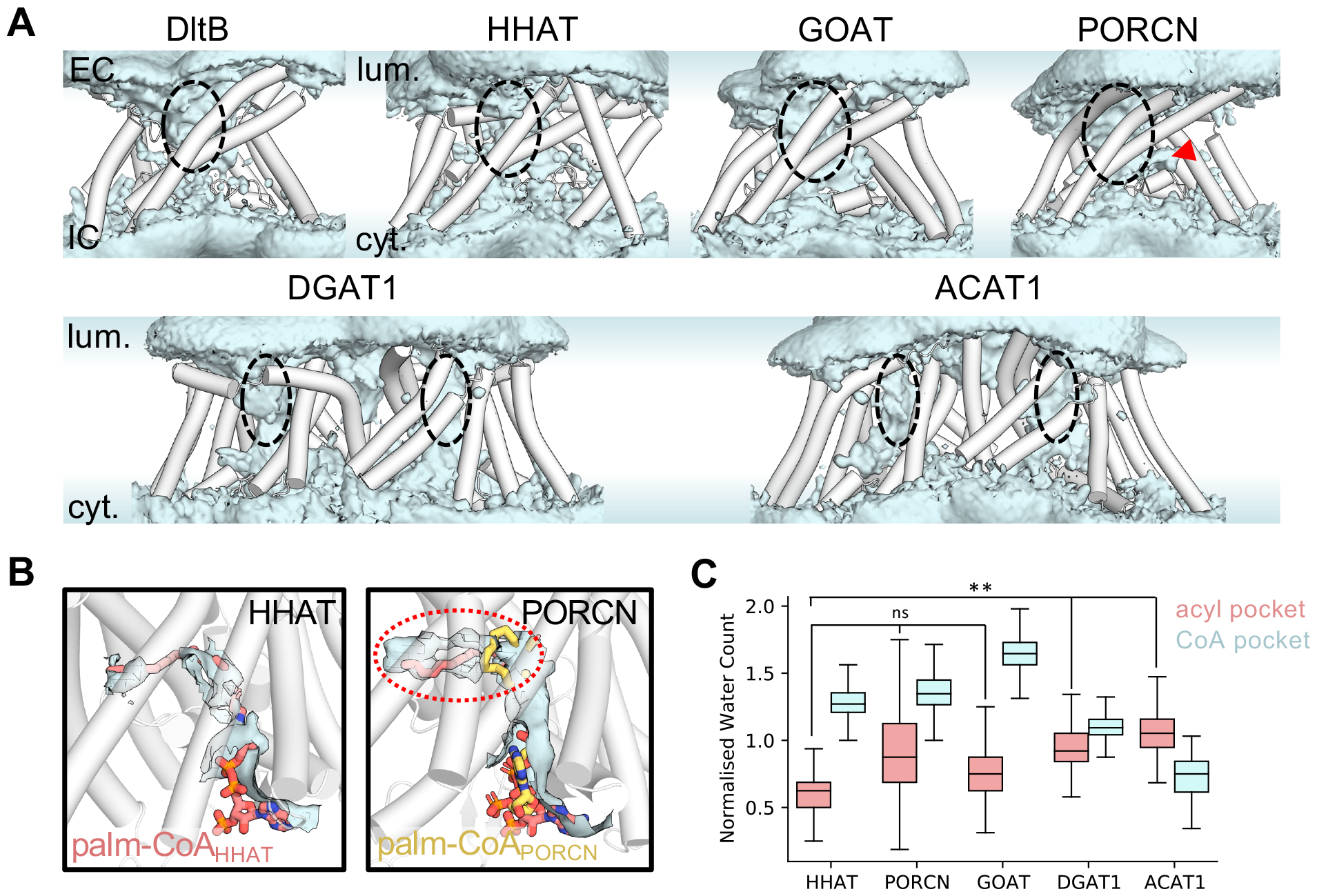
Solvation of the MBOAT reaction centre. **A)** Time averaged water density (blue isosurface) across 5 x 200 ns atomistic simulations of MBOAT enzymes. The luminal/extracellular (EC) and cytosolic/intracellular (IC) surfaces are labelled. Dashed circles indicate the location of the conserved solvated MBOAT core. The red arrow indicates the position of a hydrated projection in PORCN, shown in **B. B)** Water density (blue isosurface) in simulations of HHAT and PORCN, overlayed with the palmitoyl-CoA (salmon) and palmitoleoyl-CoA (yellow) binding conformations in structures of HHAT^9^ and PORCN^11^ respectively. The projection marked in **A** is circled. **C)** Normalised water count (see methods) within the acyl tail and CoA headgroup binding pockets across simulations of apo MBOATs. Statistical differences between lipid pocket solvation were calculated using the Students unpaired t-test: not-significant (ns): *P* > 0.05, ^*^: *P* ≤ 0.05, ^**^: *P* ≤ 0.01, ^***^: *P* ≤ 0.001, ^****^: *P* ≤ 0.0001.

Beneath the reaction centre we observe notable differences in the behaviour of water across the MBOAT family. For example, in PORCN a hydrated projection (Fig. 6A, red arrow) is present above re-entrant loop-2 in a similar position to the cavity occupied by the palmitoyl-tail of acyl-CoA in structures of HHAT^9,10^ (Fig. 6B). Hydration of this finger-like pocket is not seen in simulations of HHAT or GOAT and may explain why the palmitoleoyl-tail occupies a distinct kinked position in the PORCN structure^11^ (Fig. 6B). We further quantified solvation of the acyl-CoA binding pocket between HHAT, PORCN, GOAT, DGAT1 and ACAT1 to assess differences in the relative hydration of regions coordinating the acyl- and CoA chemical groups (see methods). For small-molecule MBOATs (DGAT1/ACAT1) the acyl pocket was more hydrated than in HHAT. By contrast the CoA coordinating pocket was less hydrated than all other protein MBOATs analysed (Fig. 6C). Hence, we observe differences in the relative hydration of regions coordinating acyl-CoA chemical groups across the MBOAT family, despite the presence of a conserved hydrated MBOAT core. These data are pertinent to the design of drugs that are a) sufficiently hydrophilic to occupy the MBOAT core, b) sufficiently hydrophobic to displace acyl-tails and c) optimised for differences in the relative hydrophobicity of distinct substrate coordinating regions for specific MBOAT targeting.

### MBOATs employ distinct mechanisms of solvent gating

Our observations of water within the MBOAT core raises the intriguing question of how hydration of the active site can occur without solvent leaking across the membrane. We previously located a hydrophobic gate within HHAT (formed by W335 and F372) which closed the palmitoyl-CoA binding cavity to the luminal accessible solvent in the absence of bound substrate, thus preventing water permeation across the membrane^9^. We identified comparable residues at equivalent positions in GOAT (S303/F331), PORCN (V302/Y334), DltB (W285/M329), DGAT1 (W374/F408) and ACAT1 (Y417/F453) and analysed atomistic simulations to assess whether they perform equivalent roles (Fig. 7).

**Figure 7:**
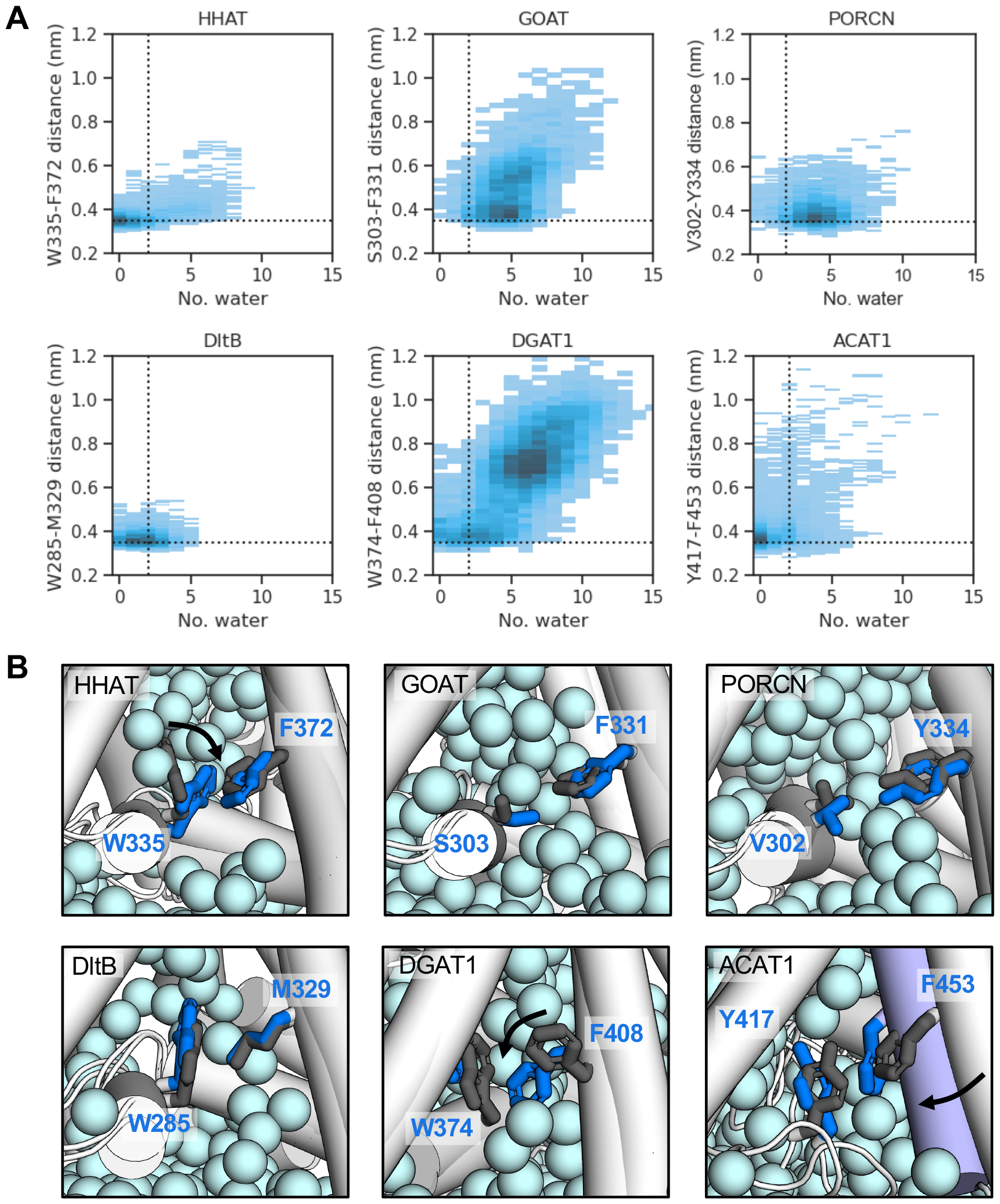
Identification of hydrophobic gating mechanisms. **A)** 2D distribution plots of the minimum distance between sidechain atoms of proposed gating residues *vs* the number of water molecules within a sphere (radius 0.4 nm) centred on the midpoint of residue pair Cα atoms. The position of the sphere was updated each frame across 5 x 200 ns simulations of MBOAT members. A dashed horizontal line at y = 0.35 nm indicates direct residue interaction. The vertical line is drawn at x = 2 waters where water permeation is prevented. **B)** Snapshots from atomistic simulations showing the position of proposed gating residue pairs at the start (grey sticks) and end (blue sticks) of trajectories. Side chain reorientations are arrowed. The oxygen atoms of water in the final snapshot are shown as light blue spheres. For ACAT1, concerted movement of TM6’ is indicated in slate.

We plotted the minimum distance between sidechain atoms of proposed gating residues *vs* the number of waters within a sphere of radius 0.4 nm centred on the Cα atoms of residue pairs (updated each frame) (Fig. 7A). Residue pairs were able to form direct interactions in all MBOATs (Fig. 7A, dashed horizontal line at 0.35 nm) however water movement was only restricted in HHAT, DltB, ACAT1 and to a lesser extent DGAT1 (Fig. 7A, dashed vertical line). In GOAT and PORCN water permeation was not prevented which we attribute to the smaller (PORCN-V302) and/or polar (GOAT-S303) residues which replace the equivalent tryptophan residues in HHAT, DltB and DGAT1 or ACAT1-Y417. Closure of the acyl-CoA pocket is permitted by movement of HHAT W335 towards F372 and DGAT1 F408 towards W374 compared to the structural conformations (Fig. 7B). DltB residue W285 and M329 remain close to the structural conformation. By contrast, the gating mechanism of ACAT1 is permitted by movement of the entire TM6’ helix towards Y417 (Fig. 7B). Hence, the location of pocket gating appears to be conserved between HHAT, DltB, DGAT1 and ACAT1 but the gating mechanism differs between side-chain flip (for HHAT, DltB and DGAT1) and concerted TM6’ movement (for ACAT1). It remains to be investigated how (and whether) GOAT/PORCN employ distinct mechanisms of protein gating given bulky hydrophobic residues are not conserved at equivalent positions in other MBOATs.

## Discussion

Cross-comparison of membrane protein features across structurally conserved families is critical for a) discerning functional specificity, b) reducing off-target effects during pharmaceutical intervention and c) tracing evolutionary lineages. Within this study we demonstrate how simulations can be effectively applied to address each of these aspects, furthering structural interpretation to encompass environmental and dynamic contexts.

Our assessment of membrane deformation suggests MBOATs differentially reduce the width of the bilayer dependant on whether they acylate proteins (40-60% reduction in bilayer width) or small-molecules (minimal perturbation) (Fig. 2, Supplementary Fig. 1-2). Thus, we define a principle and easily predictable hallmark of MBOAT reaction specificity. For protein acylating MBOATs the reduction in bilayer width may help reduce the energetic cost of reactant group (e.g. acyl) transfer across the membrane, compared to small-molecule acylating MBOATs which need only shuttle groups between the cytoplasm and the membrane^1^. Alternatively, localised regions of deformation form guiding funnels for substrate/product entry/exit away from the protein. This ‘hydrophobic steering’ proposal is conceptually similar to electrostatically driven binding of substrates to soluble enzymes to enhance catalytic rates beyond the diffusion limit^23^. Additionally, membrane thinning has been suggested to increase the diffusion rate of Rhomboid proteases, another class of membrane embedded enzymes and increase the rate of molecular collisions^24^. Intriguingly, there is a growing body of evidence in support of membrane thinning/deformation as a generalisable property of ER localised proteins which facilitate molecular transfer and protein biogenesis^25^. For example, the signal peptidase complex acts as a ‘molecular ruler’ for substrate compatibility via bilayer thinning and the DHHC acyltransferase enzyme exhibits membrane deformation towards the catalytic site^26,27^. In addition, more classical examples such as SecYEG or ERAD mediate bilayer distortion^28,29^. Hence, differential membrane thinning may be an evolutionarily conserved biophysical adaption that extends beyond the MBOAT family.

Our simulations also provide a preliminary window into the role of specific lipids in MBOAT function. For example, we observe binding of membrane derived cholesterol within the lateral gate of DGAT1 and ACAT1 (Fig. 3C). The orientation and position of cholesterol within ACAT1 matched previous docking predictions^2^ and the position of a sterol-like density in ACAT2^16^. We also observe binding of phospholipids to the luminal gate of protein acylating MBOATs, in proximity to the catalytically conserved histidine on TM6’ (Fig. 3A)^1^. During the catalytic cycle, bound cholesterol or phospholipids would presumably need to be displaced from the product exit gates during reaction cycling. Hence, designing lipid-analogues with enhanced binding affinities to these gates may represent one avenue of therapeutic inhibition. For example, lipid-like modulators have been applied to modulate ligand-gated ion channel responses^30^. Finally, we note a prolonged lipid binding pocket on DltB (Fig. 3B) which remains functionally uncharacterised. Ultimately, the role(s) of lipids in MBOATs function remains understudied, and it would be wise for future investigations to focus efforts here.

We complemented simulations with bioinformatic analyses of residue conservation across the MBOAT family. Our analyses reveal re-entrant loop-2 as a site of high conservation (across orthologues) but with key specific residue alternations across MBOAT homologues (Fig. 4). In small-molecule acylating MBOATs re-entrant loop-2 forms the dimeric interface between monomers (Supplementary Fig. 3)^2–6^. By contrast, in protein-acylating MBOATs this site is stabilised via Cys-heme-b coordination (HHAT), conserved salt-bridges (PORCN, GOAT) or *π* - *π* stacking interactions (DltB) (Fig. 4). Therefore, all MBOATs have evolved specific, conserved mechanisms to stabilise their tertiary fold at re-entrant loop-2. Each of the described molecular interactions is hypothetically reversible and hence may serve as a method of protein regulation. For example, during protein trafficking the bilayer width varies across the endosomal network^31,32^. Given the unusual MBOAT tertiary fold, it is conceivable that hydrophobic mismatch would dictate whether helices are correctly aligned for residue interactions at re-entrant loop-2, preferentially stabilising MBOATs for protein function only within their native membrane. Such bilayer thickness dependant regulation has been observed for other membrane proteins such as the Golgi localised transporter Vrg4^33^.

For DGAT1 we illuminate the role of a highly conserved hydrogen bond in DGAT1 tail-swap stabilisation (Fig. 5). This hydrogen bond between H69-T260 is uniquely conserved on the comparatively divergent cytoplasmic surface. The DGAT1 N-terminal tail has been implicated in positive cooperativity and protein regulation in previous truncation experiments^22,34^ however the molecular nature of regulation is unknown. We predict a substantial portion of this regulation may be attributed to a single, conserved hydrogen bond between H69-T260 whereby bond breaking results in dissociation of the neighbouring DGAT1 tail from the dimer.

Hence, we provide two examples of interactional divergence across the MBOAT family (surrounding re-entrant loop-2 and DGAT1 tail stabilisation) which are a) precisely localised and b) regulatorily relevant (Fig. 4-5). In addition, we describe how environmental factors external to protein structure (water, ions) must be considered when designing drugs for membrane protein targeting, including consideration of differences in the relative hydrophobicity of internal cavities (Fig. 6-7). The subtlety of solvent effects on drug binding poses is stressed further within a recent high-throughput combined simulation/experimental approach to fragment-based drug discovery^35^. These data scaffold the development of drugs for specific MBOAT targeting within a native-like context.

We end by reflecting on whether our analyses support our previously proposed hypothesis that eukaryotic MBOATs may have evolved via distinct lineages^1^. We previously noted that HHAT appears to be more closely structurally related to DltB compared to GOAT/PORCN and uniquely *N*-acylates the protein substrate in comparison to widespread *O*-acylation across the family. Furthermore, HHAT and DltB are most closely related in their mechanisms of solvent gating (Fig. 7) and HHAT is post-translationally modified at re-entrant loop-2 unlike the salt-bridge stabilisation employed by GOAT and PORCN (Fig. 4). HHAT appears to be an enigma amongst eukaryotic MBOATs. We predict that Hedgehog-mediated developmental signalling may diverge evolutionarily from biosynthetic/regulatory MBOAT lineages, ultimately affecting how functional evaluation of MBOAT mechanisms are transposed between pathways.

## Acknowledgements

T.B.A. and C.E.C acknowledge support from Wellcome (102164/Z/13/Z) while conducting this research. T.B.A. is additionally supported by Schmidt Science Fellows, in partnership with the Rhodes Trust. CEC is supported by a ResTraComp fellowship from the Hospital for Sick Children. M.S.P.S. was supported by Wellcome (208361/Z/17/Z), the BBSRC (BB/R00126X/1) and PRACE (Partnership for Advanced Computing in Europe, 2016163984). C.S. is funded by Cancer Research UK (C20724/ A26752 and DRCRPG-May23/100002), the BBSRC (BB/T01508X/1) and the European Research Council (647278).

## Author contributions

Simulations and bioinformatic analyses were performed by M.H. and T.B.A. T.B.A, C.E.C., M.S.P.S. and C.S. conceptualised the study. T.B.A. wrote the paper with input from all authors.

## Methods

### Structures and models used in simulations

Details of simulation setups are provided in Table 1. Protein coordinates for MBOAT family members DltB^13^, HHAT^9^, PORCN^11^, ACAT1^2^ and DGAT1^5^ were obtained from the Protein Data Bank (PDB), while the GOAT model was obtained from the AlphaFold Protein Structure Database^36^. The validity of the GOAT model was further assessed via conservation, electrostatics, hydrophobicity and tunnel analyses detailed in Supplementary Fig. 4. Additional proteins or ligands were removed and unresolved residues or loops were modelled using the PyMOL (https://pymol.org/2/) Mutagenesis Wizard and MODELLER9.20^39^.

**Table 1:**
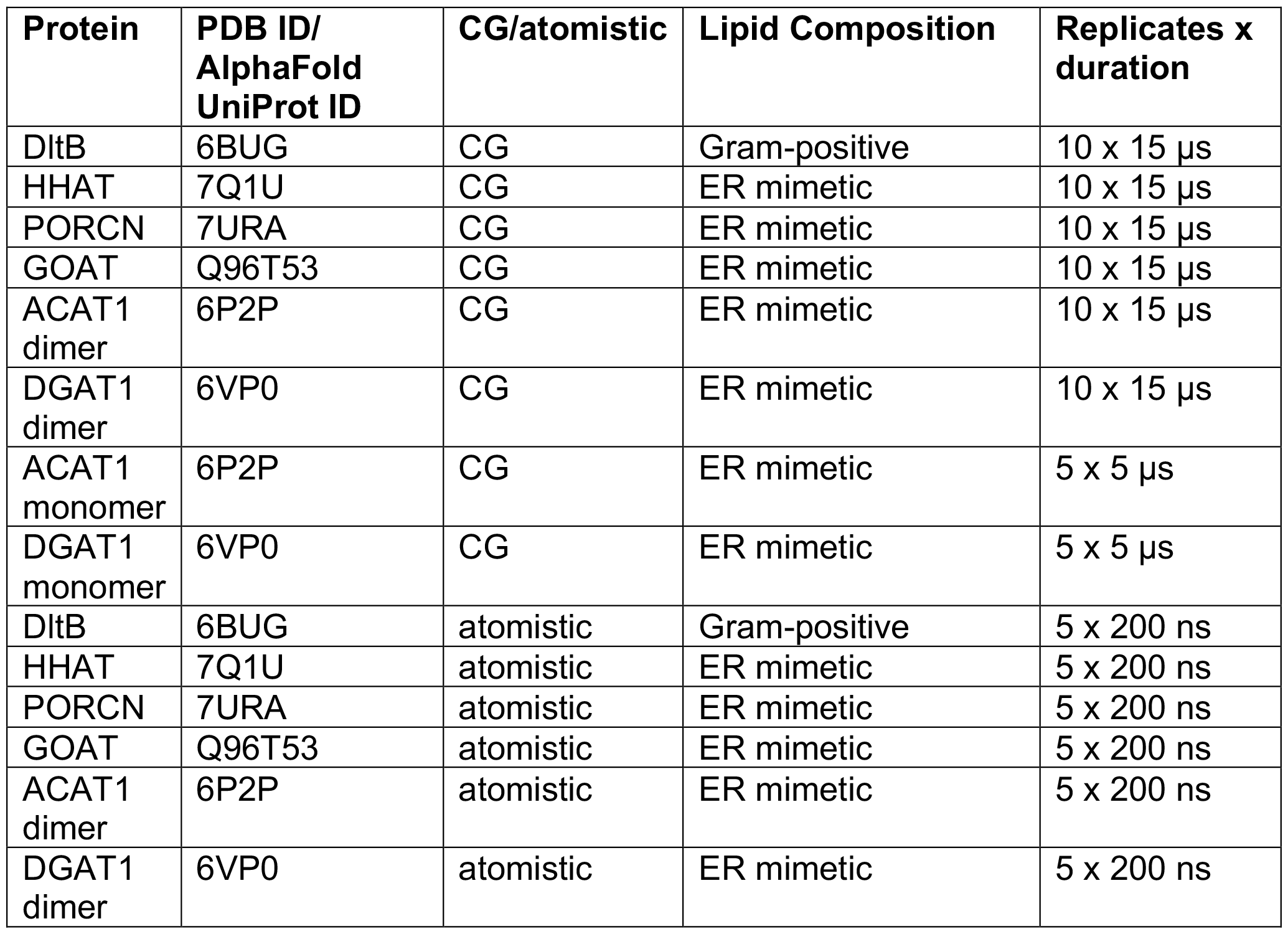
Summary of systems simulated.

### Coarse-grained MD simulations

Proteins were converted to CG resolution using *martinize*.*py* and the Martini2.2 forcefield^40,41^ with an ElNeDyn elastic network (force constant: 1000 kJ mol^-1^ nm^-2^, upper cut-off: 0.9 nm)^42^. MBOATs were embedded in bilayers designed to mimic the native membrane composition where each is localised (Fig. 1C-D). Hence, DltB was embedded in a Gram-positive like membrane composed of POPG (69%), POPE (23%) and cardiolipin (8%) and all other MBOATs were embedded in ER mimetic membranes composed of POPC (35%), DOPC (35%), POPE (8%), DOPE (7%), cholesterol (10%), palmitate (5%) in the luminal leaflet and POPC (15%), DOPC (15%), POPE (19%), DOPE (18%), POPS (8%), PIP_2_ (10%), cholesterol (10%) and palmitate (5%) in the cytoplasmic leaflet using *insane*.*py*^43^ (Table 1). The palmitate PCN Martini bead model was used and cholesterol was modelled with inclusion of virtual sites^44^. Systems were solvated using Martini water^41^ and approximately 0.15 M NaCl, in line with previously reported simulations^9^. Systems were independently energy minimised via the steepest-decent algorithm and equilibrated for 25 ns with restraints applied to all protein beads followed by a second 100 ns equilibration with restraints on protein backbone beads.

Each protein was simulated for 10 x 15 μs using the GROMACS 2019 simulation software^45^ and a 20 fs timestep. Control simulations of ACAT1/DGAT1 monomers were run for 5 x 5 μs. Temperature was maintained at 310 K using the V-rescale thermostat^46^ (*τ*_t_ = 1 ps). Pressure was maintained at 1 bar using the Parrinello-Rahman barostat^47^ (*τ*_p_ = 12 ps, compressibility = 3 x 10^−4^ bar^-1^). Periodic boundary conditions were applied. Electrostatic interactions were cut-off at 1.1 nm via the reaction-field method and van der Waals interactions were described with the potential-shift Verlet method and a 1.1 nm cut-off.

### Atomistic MD simulations

Atomistic simulations were initiated from frames backmapped from CG resolution using CG2AT^48^. Backmapping files for palmitate were not available and hence, prior to backmapping, palmitate was removed from the membrane and CG systems were re-equilibrated. The TIP3P water model^49^ was used and systems were neutralised with approximately 0.15 M NaCl. The CHARMM36 forcefield was used to describe all components^50^. Protein conformations were mapped to the structural coordinates, with corrected protonation states^51^ and modelled loops. Each system was energy minimised via the steepest decent algorithm and equilibrated in 2 x 5 ns NVT and NPT steps with restraints applied to protein heavy atoms and backbone atoms respectively.

Atomistic simulations were run for 5 x 200 ns (Table 1) with a 2 fs timestep. The GROMACS 2019 and 2020 simulation packages^45^ were used to run simulations. Long-range electrostatics were described via the Particle-Mesh-Ewald (PME)^52^ method with a 1.2 nm cut-off. Van der Waals interactions were switched from 1.0 nm to 1.2 nm using the force-switch modifier. The system was kept at 310 K and 1 bar using the Nosé-Hoover thermostat^53,54^ (*τ*_t_ = 0.5 ps) and Parrinello-Rahman barostat^47^ (*τ*_p_ = 2 ps, compressibility = 4.5 x 10^−5^ bar^-1^) respectively. A dispersion correction was not applied. Bonds were constrained to their equilibrium values using the LINCS algorithm^55^.

### Sequence conservation analysis

The MPI Bioinformatics pipeline^56^ (https://toolkit.tuebingen.mpg.de) was used for sequence conservation analysis. Uniprot sequences were used for PSI-BLAST^57^ searches with a reporting E-value cut-off of 1 x 10^−3^. Search results were inputted into the T-Coffee^58^ server to construct multiple sequence alignments (MSAs) which were subsequently mapped onto MBOAT structures using ConSurf^37^.

### Trajectory analysis

#### Membrane deformation and lipid interactions

MDAnalysis^38^ was used to calculate the z axial position of lipid phosphate beads across simulations. The mean z coordinate of all phosphate beads was taken as the bilayer midplane (z = 0 nm). Extracellular and intracellular leaflet phosphate positions were normalised to the bilayer midplane. The mean z position of phosphate beads within 0.8 nm of residues in proximity to deformations (Supplementary Fig. 1, residues marked red) were used to obtain coordinates of localised deformation towards the midplane. Global deformation was calculated as the difference between the most extreme deformations in each leaflet and all phosphate bead positions (i.e. approximately equal to at extended distances from the protein). Local deformation was defined as the difference between the most extreme deformations in each leaflet and any phosphate beads within 0.8 nm of the protein (i.e. only protein contacting phosphate beads). Specific protein-lipid interactions were calculated using PyLipID^20^ with a 0.475 nm lower and 0.7 nm upper cut-off scheme. Residence time comparison plots were adapted from LipIDens^21^.

#### Residue interactions

Analysis of salt-bridge formation was calculated using MDAnalysis with a 0.4 nm cut-off between the carboxyl O atoms of glutamate and the lysine/histidine NH groups^59^. Assessment of DGAT1 hydrogen bond formation was also performed using MDAnalysis^38^.

#### Water analyses

MDAnalysis^38^ was used to calculate the density of water O atoms within 1 nm of proteins across atomistic simulations. For calculation of solvent occupancy within the acyl-CoA binding pocket, MDAnalysis was used to align each atomistic simulation to a reference structure based on the MBOAT Cα coordinates. The reference structure corresponded to each MBOAT structure with acyl-CoA substrate bound (HHAT palmitoyl-CoA PDB: 7Q1U^9^, PORCN palmitoleoyl-CoA PDB: 7URA^11^, DGAT1 oleoyl-CoA PDB: 6VP0^5^, ACAT1 oleoyl-CoA PDB: 6P2P^2^). For the GOAT model the palmitoyl-CoA coordinates from the HHAT structure were used (Supplementary Fig. 4). Any water O atoms within 0.4 nm of atoms comprising the aligned acyl tail (up to the S atom of the thioester bond) or CoA headgroup (all other atoms) were selected for each frame. The number of water within the acyl and headgroup pockets were normalised based on the number of atoms in the acyl tail and CoA headgroup references (to account for differences in tail length and size of the chemical groups).

#### Data representation

PyMol (https://pymol.org/2/) and VMD^60^ were used for visualisation. GraphPad Prism-9 (https://www.graphpad.com) was used for calculation of statistical significance.

