## Supplementary_Information for "Mapping Structural and Dynamic Divergence Across the MBOAT family"

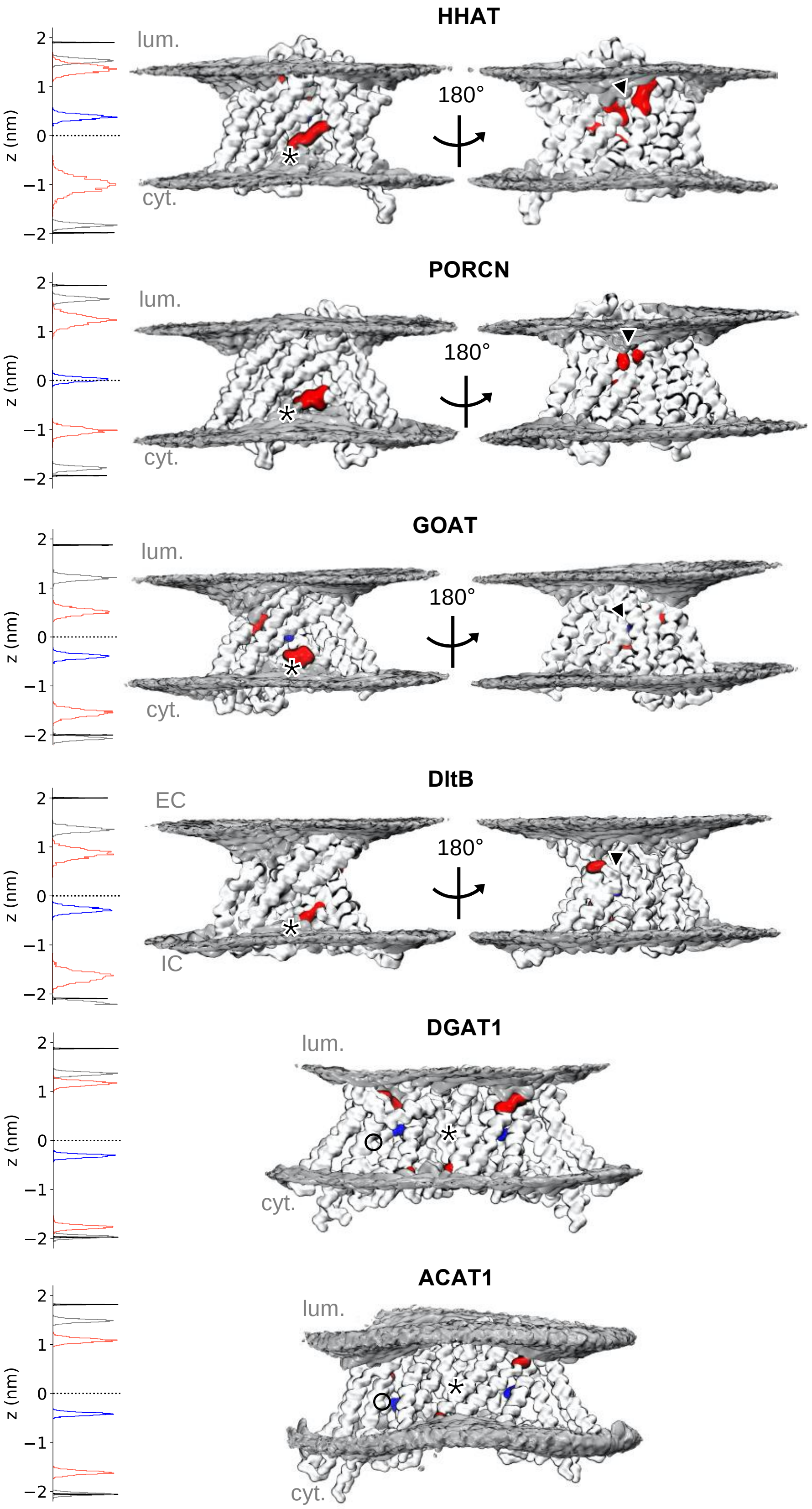
HHAT
lum.
cyt.
180°
z (nm)
PORCN
lum.
cyt.
180°
z (nm)
GOAT
lum.
cyt.
180°
z (nm)
DltB
EC
IC
180°
z (nm)
DGAT1
lum.
cyt.
z (nm)
ACAT1
lum.
cyt.
z (nm)

**Supplementary Fig. 1: Membrane deformation surrounding MBOAT family members.**

Time averaged phosphate bead density (grey volume) surrounding MBOAT family members across 10 x 15 µs CG simulations. MBOATs are coloured white in CG representation and catalytic histidines are shown in blue. Residues in proximity to the most extreme regions on deformation are coloured red. The position of luminal/extracellular (EC) and cytoplasmic/intracellular (IC) leaflets are indicated. Black asterisks mark the position of re-entrant loop-2, arrows show the location of the luminal gate and circles mark the lateral gate. Accompanying histograms show the z axial coordinates of phosphate beads within each leaflet at extended distances from the protein (black), all phosphate beads within 0.8 nm of the protein (grey) or within 0.8 nm of residues at the most extreme regions of deformation (red). The z coordinate position of the catalytic histidine backbone beads are shown in blue. Phosphate and histidine z coordinates were obtained using MDAnalysis[9] and normalised to the bilayer midplane (z = 0 nm) based on the mean position of all phosphate beads (see methods for further details).

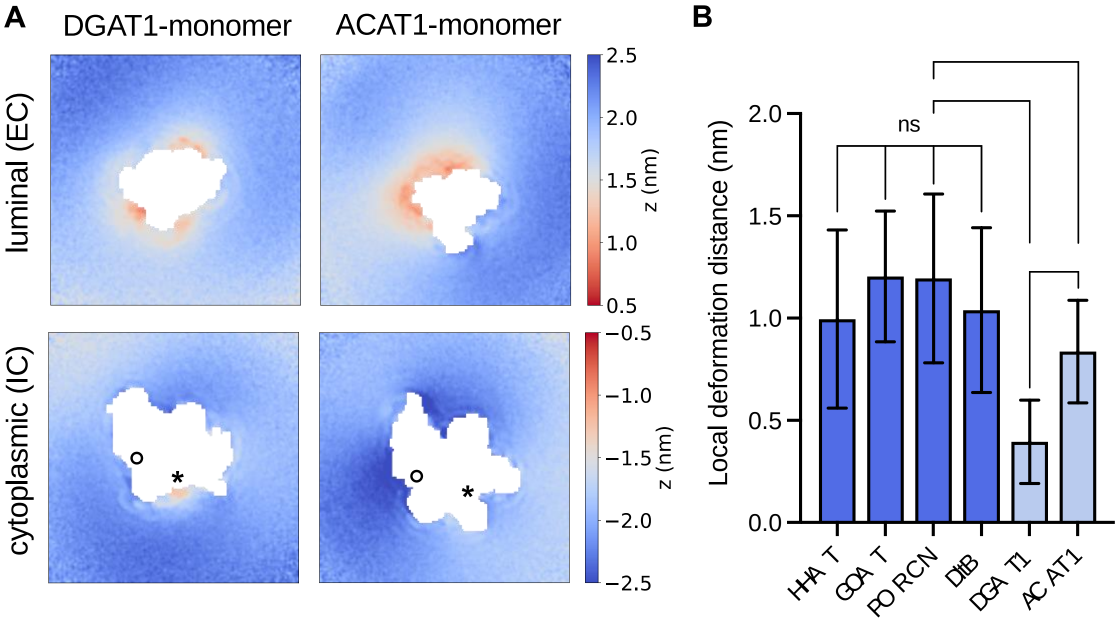


**Supplementary Fig. 2: Membrane deformation surrounding monomeric small-molecule MBOATs and localised membrane deformations across the family.**
**A)** 2D array of membrane deformation surrounding monomeric small-molecule MBOATs across control CG simulations (5 x 5 μs), defined identically to in Fig. 2. **B)** Bar plot of local membrane deformation, defined as the reduction in bilayer width between the most extreme regions of membrane deformation compared to all phosphate beads within 0.8 nm of any protein bead (i.e. protein contacting phosphates) across 10 x 15 μs CG simulations. The mean ± s.d. of phosphate bead positions is reported. Statistical significance was determined by a Students unpaired t-test: not-significant (ns): $P > 0.05$, *: $P \leq 0.05$, **: $P \leq 0.01$, ***: $P \leq 0.001$, ****: $P \leq 0.0001$.

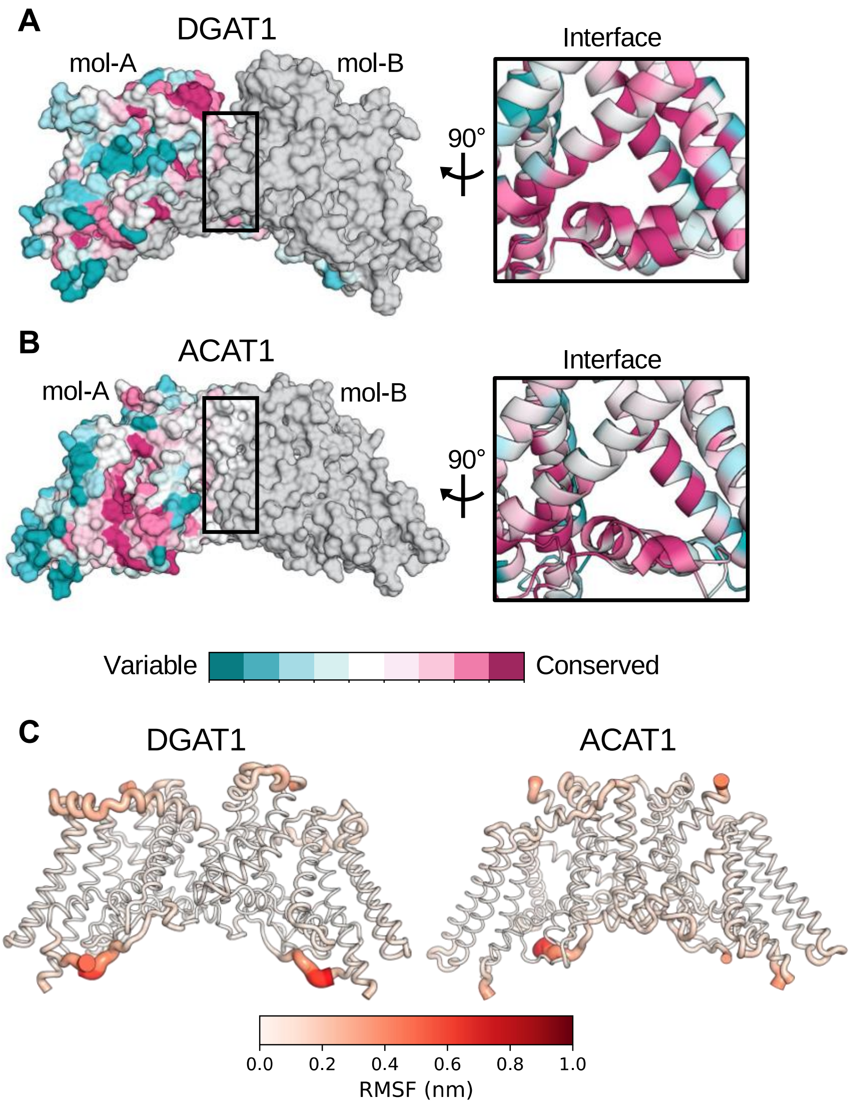


**Supplementary Fig. 3: Conservation and dynamics of small-molecule dynamics.** Per residue sequence conservation mapped onto the structures of **A)** DGAT1 and **B)** ACAT1 and coloured using ConSurf[2]**.** The second subunit within the dimer is coloured grey for clarity. The inset shows re-entrant loop-2 and surrounding transmembrane helices, as viewed from the dimeric interface (boxed). **C)** Root mean square fluctuation (RMSF) of residue Cα atoms across 5 x 200 ns atomistic simulations of DGAT1 and ACAT1 mapped onto protein structures (excluding modelled loops).

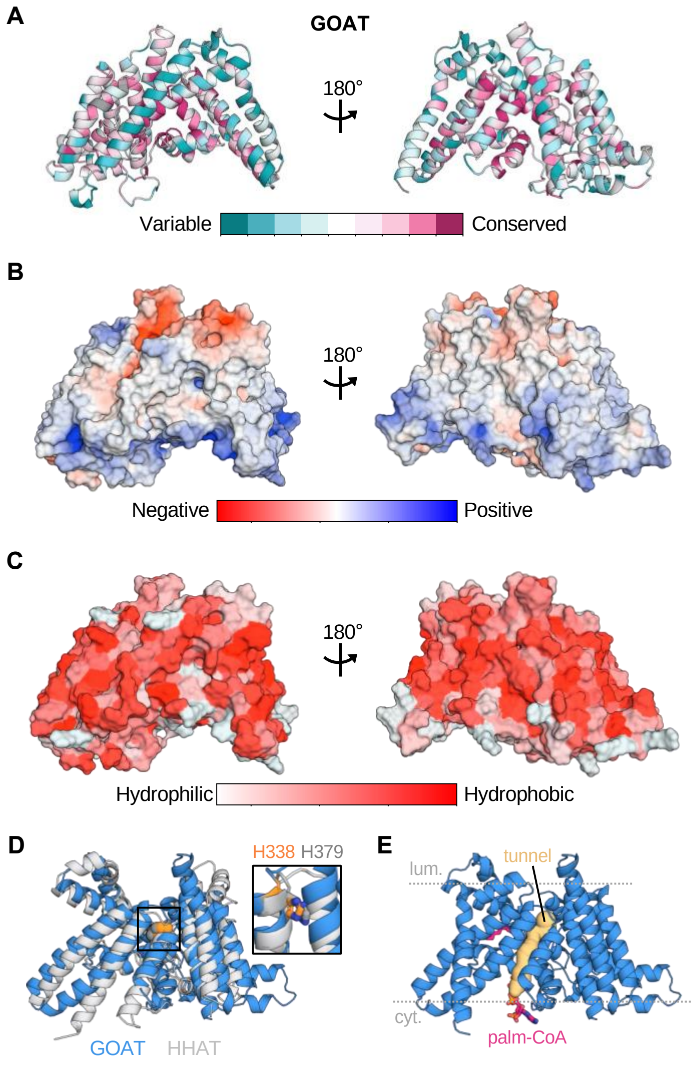


**Supplementary Fig. 4: Characterisation of the GOAT model.**
The GOAT model was obtained from the AlphaFold Protein Structure Database[1] with the UniProt ID Q96T53. The N-terminal segment with very low structural confidence prediction (residues M1-L9) was removed. **A)** Per residue sequence conservation for GOAT (see methods) mapped onto the GOAT model using ConSurf[2]. **B)** Electrostatic surface potential of GOAT obtained using the PyMol Adaptive Poisson-Boltzmann Solver (APBS) plug-in[3,4]. Charged and neutral regions align with predicted solvated and membrane exposed surfaces respectively. **C)** Distribution of hydrophobic residues on the GOAT surface coloured using the Eisenburg hydrophobicity scale[5]. **D)** Structural alignment of the GOAT (blue) and HHAT (PDB: 7Q1U[6], grey) MBOAT core helices[7]. Conserved catalytic histidines GOAT-H338 (orange) and HHAT-H379 (grey) are shown as spheres. The inset shows a close-up of the histidine overlay (stick representation). **E)** A tunnel (yellow spheres) within the predicted GOAT acyl-CoA binding pocked, obtained via the PyMol Caver3 plug-in[8]. An overlay with the palmitoyl-CoA binding pose (pink sticks) bound to HHAT[6] is shown. The position of luminal and cytoplasmic membrane leaflets are indicated by grey lines.
